# Assessment of white matter hyperintensity severity using multimodal MRI in Alzheimer’s Disease

**DOI:** 10.1101/2023.01.20.524929

**Authors:** Olivier Parent, Aurélie Bussy, Gabriel A. Devenyi, Alyssa Dai, Manuela Costantino, Stephanie Tullo, Alyssa Salaciak, Saashi A. Bedford, Sarah Farzin, Marie-Lise Béland, Vanessa Valiquette, Sylvia Villeneuve, Judes Poirier, Christine L. Tardif, Mahsa Dadar, the PREVENT-AD Research Group, M. Mallar Chakravarty

## Abstract

White matter hyperintensities (WMHs) are clinically significant MRI abnormalities often detected in the elderly and early stages of Alzheimer’s Disease. They are indicative of vascular pathology but represent a mixture of microstructural tissue alterations that is highly variable between individuals. To better understand these alterations, we leveraged the signal of different MRI contrasts sampled within WMHs, which have differential sensitivity to microstructural properties. Subsequently, we sought to examine the asso of these WMH signal measures to clinically-relevant measures such as cortical and global brain atrophy, cognitive function, diagnostic and demographic differences, and Alzheimer’s Disease-relevant cardiovascular risk factors.

Our sample of 118 subjects was composed of healthy controls (*n*=30), high-risk of Alzheimer’s Disease due to familial history (*n*=47), mild cognitive impairment (*n*=32), and clinical Alzheimer’s Disease (*n*=9) as a means of ascertaining a spectrum of impairment. We sampled the median signal within WMHs on weighted MRI images that are commonly acquired (T1-weighted [T1w], T2-weighted [T2w], T1w/T2w ratio, Fluid-Attenuated Inversion Recovery [FLAIR]), and the relaxation times from quantitative T1 (qT1) and T2* (qT2*) images. Main analyses were performed with a periventricular/deep/superficial white matter parcellation and were repeated with a lobar white matter parcellation.

We demonstrated that the correlations between WMH signal measures were variable, suggesting that they are likely influenced by different microstructural properties. We observed that the WMH qT2* and FLAIR measures displayed different age- and disease-related trends compared to normal-appearing white matter, highlighting sensitivity to WMH-specific tissue deterioration. Further, WMH qT2* particularly in periventricular and occipital white matter regions was consistently associated with several of our clinical variables of interest using both parcellation schemes in univariate analyses, and further showed high contributions to a pattern of brain variables that was associated with age and cognitive variables in multivariate Partial Least Squares Correlation analyses. qT1 and FLAIR measures showed consistent clinical relationships in multivariate analyses only, while T1w, T2w, and T1w/T2w ratio measures were not consistently associated with clinical variables.

We observed that the qT2* signal was sensitive to clinically-relevant microstructural tissue alterations specific to WMHs. Combining volumetric and signal measures of WMH, particularly qT2* and to a lesser extent qT1 and FLAIR, should be considered to more precisely characterize the severity of WMHs in vivo. These findings may have implications in determining the reversibility of WMHs and potential efficacy of cardio- and cerebrovascular treatments.

## 1. Introduction

White matter hyperintensities (WMHs) are areas of higher magnetic resonance imaging (MRI) signal within white matter on T2-weighted (T2w) and Fluid Attenuated Inversion Recovery (FLAIR) images. WMHs are commonly detected in the elderly and considered to be markers of small vessel disease. The vascular disease burden, often measured with the total WMH volume, is increasingly recognized to play an important role in cognitive manifestations of Alzheimer’s Disease (AD),^1–3^ with abnormalities in WMH volume detected up to 20 years before AD diagnosis.^4^ Increased WMH volumes have also been consistently associated with impaired cognition in otherwise cognitively normal individuals,^5–7^ cortical atrophy,^8–11^ and cardiovascular risk factors.^12–14^

However, ex vivo histological examinations of microstructural alterations within WMHs report heterogeneous tissue alterations,^15,16^ with demyelination, axonal loss, and inflammation being present at various degrees or even absent.^17–20^ There is a crucial need to better assess the severity of WMH microstructural alterations in vivo. This could be attained by leveraging MRI signal information within WMHs on commonly-acquired normalized T1w, T2w, T1w/T2w ratio (a non-specific proxy for myelin concentration) and FLAIR images (where the hyperintense appearance of WMH is typically observed), or by assessing specific biophysical tissue properties of WMHs via novel quantitative MRI acquisitions directly measuring T1 and T2* relaxation times. In white matter, quantitative T2* relaxation time (qT2*; the sum of R2 and magnetic susceptibilities) has been linked to iron, myelin, and fiber orientation,^21,22^ whereas the quantitative T1 relaxation time (qT1) is influenced by myelin to a larger extent than iron.^22^ Quantitative MRI indices are minimally impacted by non-biological factors, such as field inhomogeneities, scanner hardware, and sequence parameters relative to weighted MRI techniques.^23^ Despite these advantages, quantitative MRI is sparingly used in clinical practice. In this study, we aimed to determine which WMH microstructural measures would be useful in assessing the severity of tissue damage. We measured six different MRI signals within WMHs in a sample of cognitively intact elderly and of participants spanning the AD spectrum (from high-risk to clinically diagnosed AD dementia) totaling 118 subjects. Using univariate and multivariate methods and two different white matter parcellations, we examined the relationships among WMH measures and assessed the sensitivity of signal trends to WMH-specific tissue deterioration by comparing the age- and disease-related trends of the signal between WMH and normal-appearing white matter (NAWM). We further related WMH measures to multiple types of clinical variables (cortical and global atrophy, cognition, demographic and group differences, and cardiovascular risk factors), with the rationale that signal measures sensitive to clinically-meaningful variations in underlying WMH tissue alterations would be related to adverse neurobiological and clinical outcomes.

## 2. Materials and methods

### 2.1 Participants

Our methodology is outlined in Fig. 1. The data was acquired as part of the Alzheimer’s Disease Biomarkers (ADB)^24–27^ and PREVENT-AD^28,29^ cohorts. Signed informed consent from all participants was obtained and the research protocols were approved by the Research Ethics Board of the Douglas Mental Health University Institute, Montreal, Canada. Summary statistics of demographic and cognitive variables before and after quality control (QC) procedures (see Supplementary Methods 1) are detailed in Table 1 (both cohorts combined). ANOVAs and Chi-Squared tests were used to assess differences in the samples before and after QC for continuous and categorical variables respectively. No significant differences were observed at the *p*<0.05 level.

**Table 1.** Summary statistics of demographic and cognitive variables before and after quality control (QC) procedures.

| Variable | Before quality control |  | After quality control |  | P-values |
| --- | --- | --- | --- | --- | --- |
|  | Statistic | Missing values | Statistic | Missing values |  |
| Total N | 253 | - | 118 | - | - |
| Group (HC, FAMHX, MCI, AD) | 82, 78, 63, 30 | 0 | 30, 47, 32, 9 | 0 | $p = 0.1902$ |
| Age | 68.0 +/- 6.69<br>(54-91) | 1 | 66.6 +/- 6.33 (54-83) | 0 | $p = 0.0531$ |
| Sex (male, female) | 96, 157 | 0 | 42, 76 | 0 | $p = 0.7481$ |
| BMI | 26.5 +/- 4.68 | 1 | 26.1 +/- 4.26 | 0 | $p = 0.4235$ |
| Years of education | 15.0 +/- 8.08 | 5 | 15.8 +/- 3.96 | 0 | $p = 0.3187$ |
| Number of APOE-ε4 alleles (0, 1, 2) | 156, 87, 10 | 0 | 75, 37, 6 | 0 | $p = 0.7761$ |
| AD8 | 1.2 +/- 1.76 | 11 | 1.1 +/- 1.66 | 2 | $p = 0.8080$ |
| MoCA | 24.7 +/- 4.42 | 5 | 25.1 +/- 3.94 | 1 | $p = 0.4015$ |
| RBANS attention | 98.9 +/- 15.76 | 30 | 100.3 +/- 14.06 | 10 | $p = 0.6428$ |
| RBANS delayed memory | 91.92 +/- 18.19 | 33 | 91.8 +/- 17.71 | 10 | $p = 0.5625$ |
| RBANS immediate memory | 93.9 +/- 17.55 | 31 | 94.6 +/- 16.79 | 10 | $p = 0.8791$ |
| RBANS visuospatial memory | 95.1 +/- 15.64 | 31 | 94.9 +/- 15.74 | 10 | p=0.6226 |
| RBANS language | 97.1 +/- 12.55 | 29 | 97.6 +/- 14.35 | 9 | p=0.8546 |
| RBANS total | 93.8 +/- 14.68 | 33 | 94.4 +/- 14.63 | 10 | p=0.7784 |
| Alcohol (yes, no) | 204, 41 | 8 | 103, 12 | 3 | p=0.1335 |
| Diabetes (yes, no) | 17, 227 | 9 | 8, 107 | 3 | p=1 |
| High BP (yes, no) | 76, 169 | 8 | 31, 84 | 3 | p=0.4198 |
| High cholesterol (yes, no) | 86, 158 | 9 | 38, 77 | 3 | p=0.6501 |
| Smoking (yes, no) | 17, 227 | 9 | 7, 108 | 3 | p=0.8842 |

Mean +/- standard deviations are shown for continuous variables, frequencies for categorical variables, and the age range is displayed. The number of missing values for each variable was computed. The column of missing values after QC also refers to the number of computationally imputed observations included in the analyses. ANOVAs and Chi-Squared tests were used to assess differences in the samples before and after QC for continuous and categorical variables respectively. P-values of those tests are shown. (QC: quality control; HC: healthy controls; FAMHX: familial history of Alzheimer's Disease; MCI: mild cognitive impairment; AD: Alzheimer's Disease dementia; BMI: body-mass index; MoCA: Montreal Cognitive Assessment; RBANS: Repeatable Battery for the Assessment of Neuropsychological Status; BP: blood pressure)

**Figure 1.**
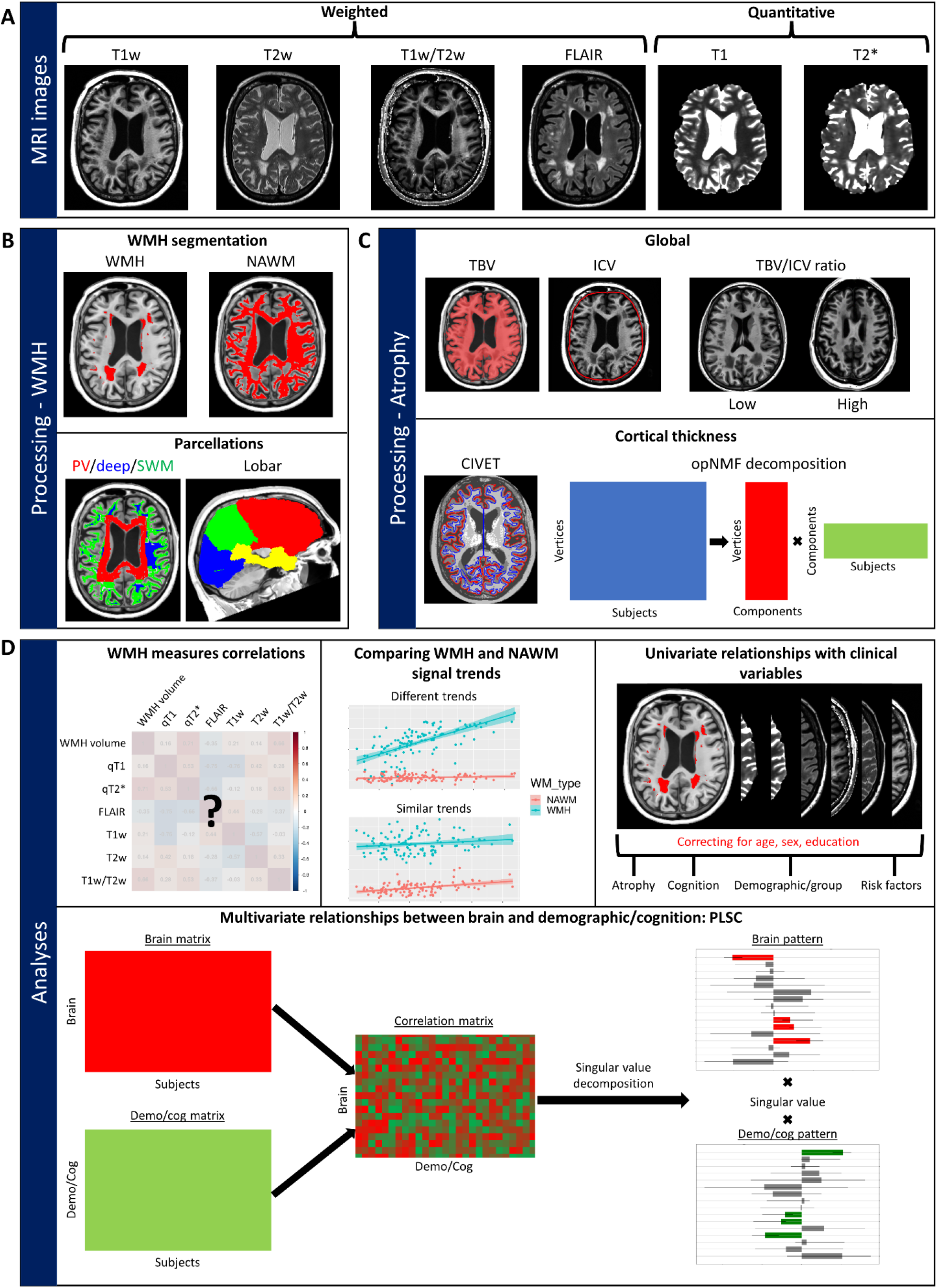
Workflow of MRI acquisitions, processing and analyses. (**A**) Six types of MR images were acquired and processed. Figures are from a 74 year old female participant with AD and high WMH burden. (**B**) Top: White matter tissue was separated into WMH and NAWM; Bottom: both tissue types were parcellated with a periventricular (PV)/deep/superficial white matter (SWM) and lobar parcellation. Subject-wise median signal was sampled within each subregion and MRI image. (**C**) Atrophy measures included global (total brain volume [TBV], intracranial brain volume [ICV], TBV/ICV ratio) (top) and cortical thickness (CT) measures (bottom). The dimensionality of the vertex-wise CT data was reduced by deriving a data-driven parcellation using non-negative matrix factorization (decomposition process is shown). (**D**) Four types of analysis were performed: 1) correlations of WMH signal measures between themselves, 2) comparing WMH and NAWM signal age- and disease-related trends, 3) assessing the univariate relationships of WMH measures with clinically-relevant variables (atrophy, cognition, clinical group, cardiovascular risk factors), and 4) investigating multivariate relationships between brain and demographic and cognitive variables with Partial Least Squares Correlation (PLSC).

### 2.2 Data acquisition

#### 2.2.1 Demographics, cognition, and cardiovascular risk factors

Participants were recruited across the AD spectrum and labeled as one of four different groups: healthy controls (HC), high-risk due to familial history of Alzheimer’s Disease (FAMHX), mild cognitive impairment (MCI), and Alzheimer’s Disease (AD) dementia. MCI and AD participants were referred to this study after diagnosis by the clinical team at the McGill Centre for Studies in Aging in Montreal, Canada. FAMHX participants were recruited by the PREVENT-AD group and had a Clinical Dementia Rating of 0 and had at least one parent diagnosed with AD. HC participants were recruited through advertisements in local newspapers targeted towards aging populations, Facebook, and Kijiji posts. Exclusion criteria included psychiatric and intellectual disorders, brain damage and concussion, current use of psychoactive substances, and contraindications to MRI.

Demographic variables included the body-mass index (BMI) and the number of APOE-ε4 alleles calculated with the PCR method^30^ using the Pyrosequencing protocol recommended by the manufacturer. Cognitive assessments included the AD8,^31^ the Montreal Cognitive Assessment (MoCA),^32^ and the Repeatable Battery for the Assessment of Neuropsychological Status (RBANS).^33^ We acquired the self-reported history of alcohol consumption, diabetes, high blood-pressure (BP), high cholesterol, and smoking in a binary format (yes or no).

Missing values for demographic, cognitive, and risk factor variables (see Table 1) were imputed with Random Forest imputation using the missForest version 1.5 package in R^34^ on the complete sample before exclusions based on MRI quality control.

#### 2.2.2 MRI acquisition

Identical MRI sequences were acquired for both cohorts (ADB and PREVENT-AD) on a Siemens Trio 3T scanner using a 32-channel head coil at the Cerebral Imaging Center, associated with the Douglas Research Center in Montreal, Canada. Each resulting image for one subject is visualized in Fig. 1A. Acquisition parameters of MRI protocols, including T1w, T2w, FLAIR, quantitative T1 (from an MP2RAGE sequence), and T2* (from a 12-echo GRE sequence) images are detailed in Supplementary Methods 2. All images are 1 mm isotropic or of higher resolution.

### 2.3 Image Processing

#### 2.3.1 Global atrophy measures

Prior to brain segmentation, we preprocessed the T1w images with minc-bpipe (https://github.com/CobraLab/minc-bpipe-library), which performed N4 field inhomogeneity correction,^35^ cropping of the neck region, and brain mask extraction using the BEaST nonlocal segmentation technique.^36^ Broad tissue types were segmented with the MINC Classify tool (https://github.com/BIC-MNI/classify). Total brain volume (TBV) was calculated as the total volume of gray and white matter in mm^3^. Intracranial brain volume (ICV) was calculated as the determinant of the transformation (i.e., a volume scaling) of a skull-to-skull registration of the T1w image to the MNI ICBM152 template. We further combined these two metrics by calculating the ratio of TBV to ICV (TBV/ICV ratio), representing a global measure proportional to the emptiness inside the skull, and thus representing global atrophy (Fig. 1B).

#### 2.3.2 Cortical thickness

We used the CIVET 2.1.0 pipeline to generate cortical surfaces (Fig. 1B).^37,38^ Cortical thickness was estimated at each vertex as the Laplacian distance between the gray-white matter boundary surface and the pial surface. Values were then resampled into MNI space and surface smoothed with a 20 mm full-width half-max heat kernel.^39^

To reduce the dimensionality of cortical thickness data, we derived a data-driven parcellation using orthogonal projective Non-Negative Matrix Factorization (NMF),^40–42^ further detailed in Supplementary Methods 3. Briefly, this method identifies covariance patterns by deconstructing an input matrix of vertices by subjects into two matrices: 1) vertices by components (representing the spatial parcellation) and 2) components by subjects (proportional to the cortical thickness of every subject inside each component) (Fig. 1B). The second matrix is used as the subject-wise measure of cortical thickness. The number of components is determined by analyzing the stability and accuracy of the reconstruction across different component granularities.

#### 2.3.3 T1w/T2w ratio processing

The T1w/T2w ratio developed by Glasser et al.^43^ has been proposed as being more myelin-sensitive than either contrast alone. To generate the T1w/T2w ratio images, we first downsampled the T2w images from 0.64 mm to 1 mm isotropic and rigidly registered the T2w images to the subject-specific T1w images (to have matched resolution and space). We then divided the raw T1w images by the matched T2w images, as in Tullo *et al*.^*26*^

#### 2.3.4 White matter hyperintensity segmentation

Before WMH segmentation, we rigidly registered T2w and FLAIR images to the subject-specific T1w images with the Advanced Normalization Tools (ANTs) pipeline^44^ and preprocessed all images by performing denoising,^45^ N3 field inhomogeneity correction,^46^ and intensity normalization. Non-linear registration of the ADNI template to the subject-specific T1w image was performed with ANTs.^44^ WMHs were segmented using a validated and automated random forest classifier.^47,48^ We used the preprocessed T1w and T2w images as inputs. Despite being classically used for WMH segmentation, FLAIR images were not included in our WMH segmentation processing because the contrast of gray to white matter of our FLAIR images was different to the contrast of the training data, which led to a higher rate of false positive WMH segmentations near the cortical gray matter. As a result, it is possible that less severe lesions were under-segmented. NAWM masks were created by removing the WMH mask dilated by 2 mm from the global white matter mask (Fig. 1B).

#### 2.3.5 White matter parcellations

While the classical parcellation of WMH only segregates periventricular (PV) and deep white matter regions, we further differentiated WMHs located in the superficial white matter (SWM) (Fig. 1B). The PV mask was obtained by dilating a mask of the ventricles by 8 mm, similarly to other studies.^49–52^ The SWM mask was obtained by dilating the cortical gray matter mask by 1 mm. In cases where WMHs were in both PV and SWM masks, the WMH was classified as PV. The deep white matter mask contained the remaining white matter voxels. WMH lobar localization has been shown to differentially relate to cognition and dementia.^53^ To estimate lobe-specific WMH burden, we used non-linear registration to map the Hammers atlas^54–56^ from ADNI to native T1w space (Fig. 1B).

### 2.4 White matter hyperintensity measures

In each white matter region (i.e., global and parcellated), we derived the WMH volumes as well as six WMH signal measures: T1w, T2w, T1w/T2w ratio, FLAIR, qT1, qT2*. The WMH volume was calculated in mm^3^, divided by TBV, and log-transformed, as recommended in previous studies.^6^ Signal measures were derived by calculating the median signal in each region for each MRI contrast. For quantitative images, we sampled the raw intensities. For qualitative images, we sampled intensities on the bias-field corrected images which we further normalized by dividing by the image-specific median intensity in the genu of the corpus callosum (defined by a manually segmented mask registered to native space), a region where no WMHs were observed in our sample, to limit non-biological sources of between-subject intensity differences. A low number of subjects had less than five WMH voxels in some white matter regions (12 subjects without deep WMHs, and 2 subjects without SWM WMHs). Instead of removing those subjects, which would bias our sample towards subjects with more advanced pathology, we imputed WMH volume values by log-transforming 1/TBV since it is impossible to divide 0, and WMH signal values by the NAWM signal value in the same region for each MRI image (e.g., qT1 in deep WMH was replaced by qT1 in deep NAWM).

### 2.5 Statistical analysis

All univariate analyses were performed with R/3.5.1. The whole sample, with all clinical groups combined, was analyzed together in order to leverage the full variability of WMH severity, neurodegeneration, and cognitive functioning across the AD spectrum, resulting in higher statistical power to detect clinical associations. First, to determine if the WMH signal measures were redundant or if they were each sensitive to different sources of microstructural variations, we computed cross-correlations matrices (*p*<0.01 threshold) between the WMH characteristics in a within-region between-measure fashion, and in a between-region within-measure fashion (Fig. 1C).

Second, we sought to assess if the WMH signal trends simply represented deterioration of the global white matter, or if these trends were specific to the lesions. We thus calculated the divergence of slopes between WMH and NAWM signal relative to age and WMH volume, which we interpreted as indicators of time and vascular burden, respectively. We used linear models with an interaction term (Eq. 1-2) and thresholded at the *p*<0.01 level.

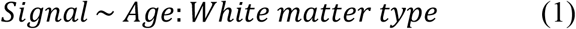

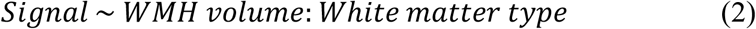

Third, to assess the clinical relevance of the microstructural variation captured by the different WMH signal measures, we used linear models to individually relate WMH measures to four types of clinical variables: cortical and global atrophy, cognition, demographic (including clinical groups), and cardiovascular risk factors (Fig. 1C), while covarying for age, sex, and years of education. We further assessed the added predictive value above the WMH volume for each WMH signal measure by additionally covarying for the region-specific WMH volume (e.g., for PV qT2*, we added PV WMH volume as a covariate). All continuous variables were z-scored before analysis to obtain standardized beta coefficients. We then examined the p-values of the clinical term thresholded at *p*<0.01. We further corrected p-values within the relationship matrix with False Rate Discovery (FDR) correction.^57^ Relationships that survived FDR correction at the 0.1 level were reported. For all significant group effects, pairwise differences were investigated post-hoc with Tukey contrasts. The standardized beta coefficients, 95% confidence intervals, p-values, and FDR-corrected p-values for all univariate relationships are available on our GitHub (https://github.com/CoBrALab/WMH_Signal_AD_OParent_2023).

Fourth, in order to investigate how signal measures of WMH are related to demographics and cognition in conjunction with traditional brain markers (i.e., cortical thickness and WMH volume), we used the multivariate technique Partial Least Squares Correlation (PLSC), detailed in Supplementary Methods 4. Briefly, we used behavioral PLSC with the Python/3.9.7 package pyls/0.01 (https://github.com/rmarkello/pyls), which performed singular value decomposition on a correlation matrix that relates each brain variable to each cognitive and demographic variable (Fig. 1C).^58–61^ This results in latent variables (LV) representing linear combinations of cognitive and demographic variables that maximally covary with linear combinations of brain variables. We inverted the directionality of LVs when appropriate for ease of interpretation. Of note, diabetes and smoking history variables were discarded from this analysis since they did not have the required level of variance for PLSC.

### 2.6 Data availability

All processing and analysis software used are free and open-access. The final data matrix, analysis code, and results in raw form are available on our GitHub page (https://github.com/CoBrALab/WMH_Signal_AD_OParent_2023).

## 3. Results

### 3.1 White matter parcellations

Results of white matter parcellations are shown in Supplementary Fig. 1. In the PV/deep/SWM parcellation, the vast majority of WMH voxels were classified as PV (86%), relatively few were classified as SWM (11%), and a very small fraction were classified as deep (2%). Importantly, the median number of voxels included in deep WMHs was only 20, potentially rendering the calculation of the median signal inside these regions less reliable and more influenced by outliers. In the lobar parcellation of WMHs (Supplementary Fig. 1B), the most affected region was frontal (58%), followed by parietal (17%), temporal (13%), and occipital (8%).

### 3.2 Spatially varying and moderate correlations between WMH measures

The correlations of WMH measures were analyzed within-region between-measure (Fig. 2A) and between-region within-measure (Fig. 2B) with correlations thresholded at *p*<0.01. Global and PV between-measure relationships were highly similar and generally showed low to moderate correlations (*r*<0.6) with some exceptions of high correlations (i.e., WMH volume with qT2* and FLAIR; qT1 with T1w). Compared to PV regions, SWM had higher correlations between qT2*, qT1, and FLAIR, while in deep white matter, correlations were higher between WMH volume, T1w, T2w, and T1w/T2w ratio.

**Figure 2.**
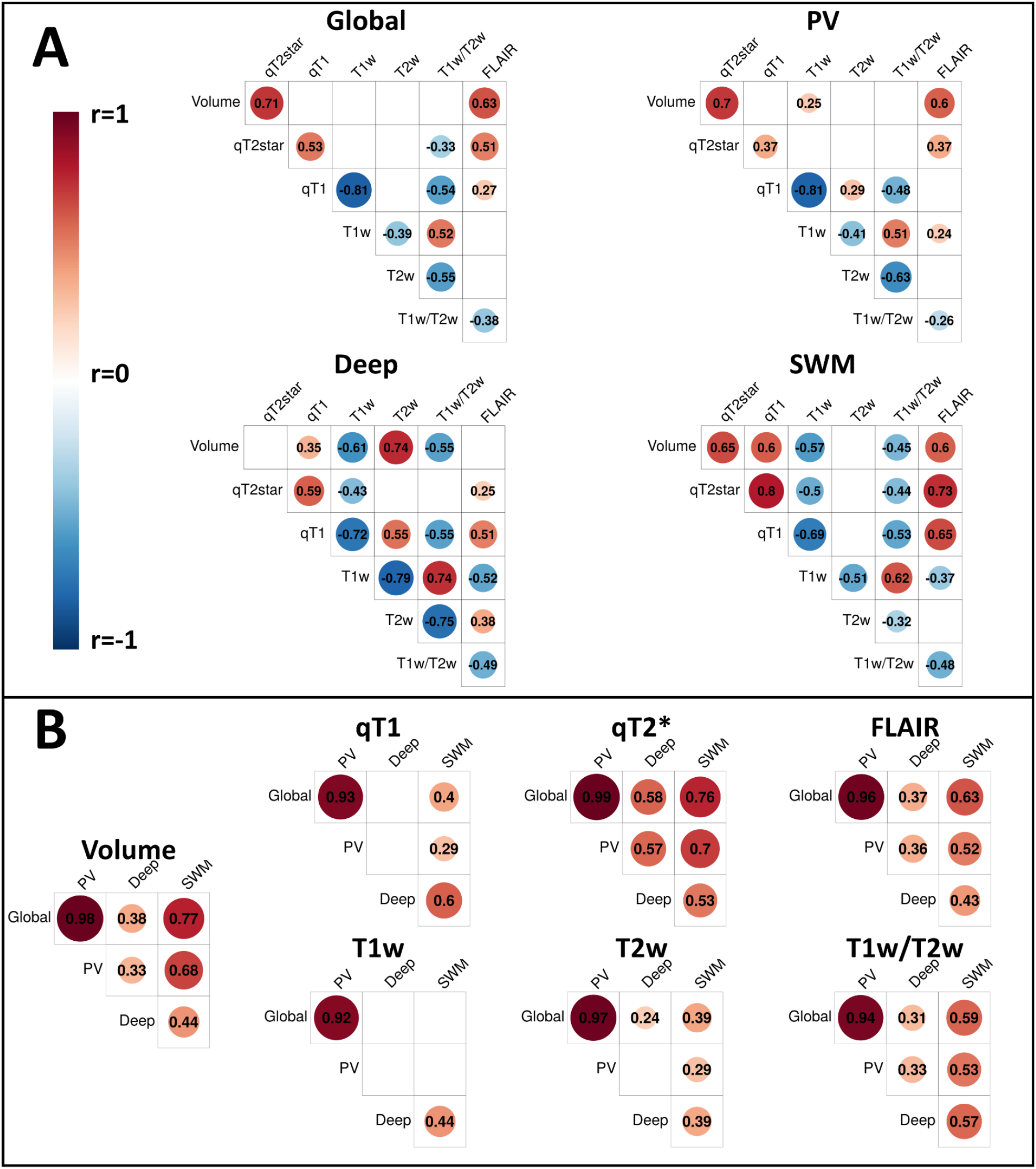
Correlations of WMH measures. (**A**) Within-region between-measure correlations of all WMH characteristics (volume and signal measures). (**B**) Between-region within-measure correlations. Circles are proportional to the amplitude of the correlation coefficients, which are also indicated. Warmer colors indicate positive correlations, and colder colors indicate negative correlations. Only significant correlations at *p*<0.01. are displayed

Subsequently, between-region within-measure relationships were analyzed (Fig. 2B). For all WMH characteristics (volume and signal measures), global and PV regions were very highly correlated (*r*>0.9). For WMH volume, qT2*, and FLAIR measures, correlations were highest between global, PV, and SWM regions, and deep regions showed lower correlations with other regions. An inverse pattern was observed for qT1 and T1w showing higher relationships between deep and SWM regions compared to other region pairs. T2w and T1w/T2w ratio measures showed higher correlations between SWM and other regions.

### 3.3 qT2* and FLAIR are sensitive to WMH-specific tissue degradation

We investigated if the WMH signal trends simply represented deterioration of the global white matter, or if these trends were specific to the lesions. We calculated the divergence of the slopes between WMH and NAWM relative to age and WMH volume (Fig. 3A). Results showed significantly different age- and disease-related signal trends between WMH and NAWM for FLAIR and qT2* in every region except deep white matter. For FLAIR and qT2*, WMH signal increases with age and WMH volume, while NAWM signal remains relatively constant (Fig. 3B). For other measures, there is a large baseline difference between NAWM and WMH but signal trends are highly similar. All significant effects at *p*<0.01 also survived FDR correction at the 0.05 level.

**Figure 3.**
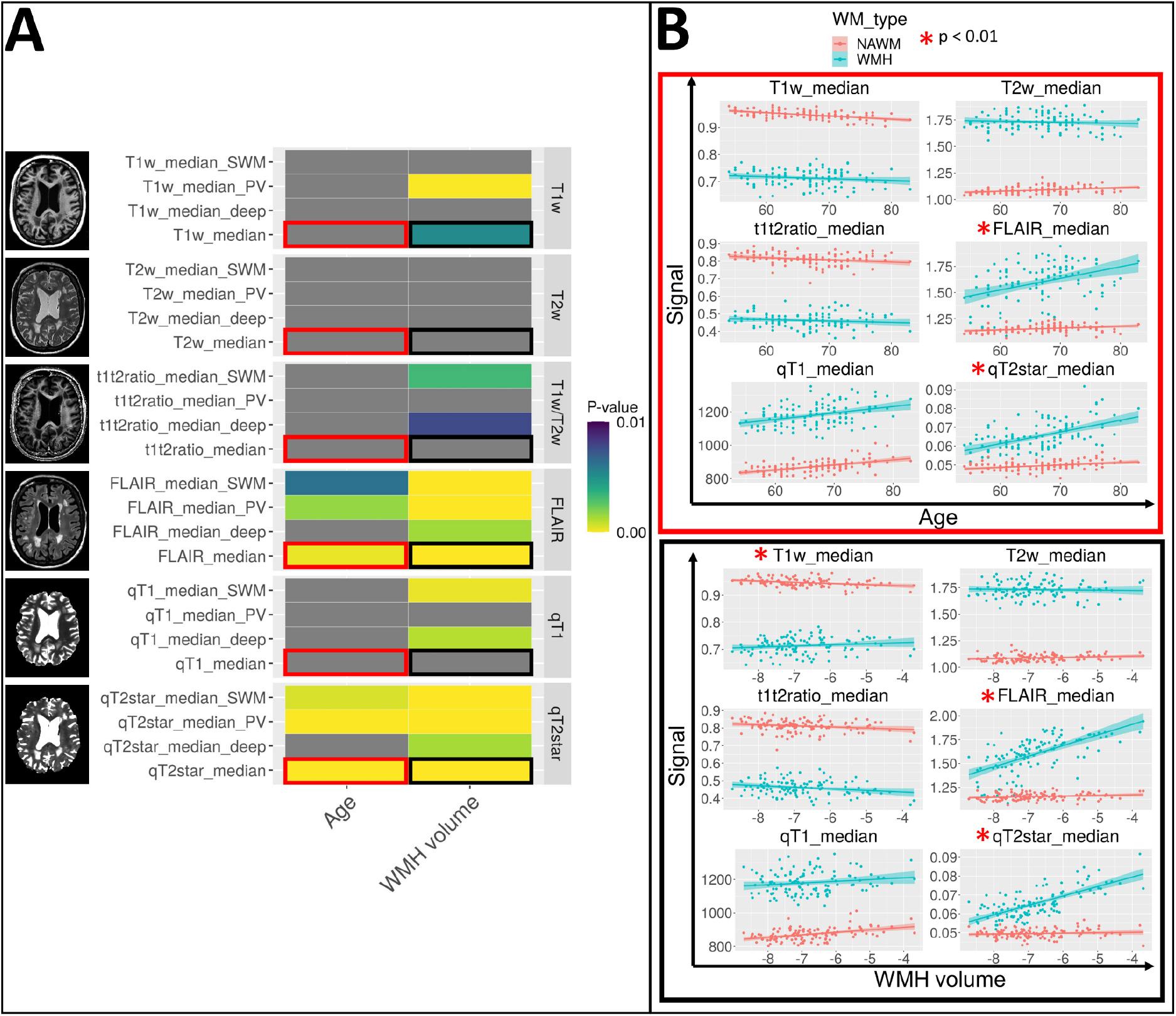
Comparing signal trends between WMH and NAWM. (**A**) On the y-axis, all WMH signal measures (global and parcellated) are grouped by image type. P-values for the interaction between either age (left) and WMH volume (right) with white matter type thresholded at *p*<0.01. are color coded, and non-significant associations are in gray. Yellow colors indicate lower p-values, and blue colors indicate higher p-values. Red squares indicate relationships for the age term that are visualized, and black squares indicate relationships for the WMH volume term that are visualized. (**B**) Graphical visualization of NAWM (red) and WMH (blue) signal trends in global white matter with age (top) and WMH volume (bottom) for each image type. Significantly different white matter trends at the *p*<0.01. level are indicated with a red star.

### 3.4 Data-driven cortical thickness parcellation

To reduce the dimensionality of the cortical thickness data, the NMF technique was used to derive a data-driven parcellation of cortical thickness. We chose to use eight components since the stability plateaued at 6 components, while the accuracy increased substantially up to eight components (Fig. 4).

**Figure 4.**
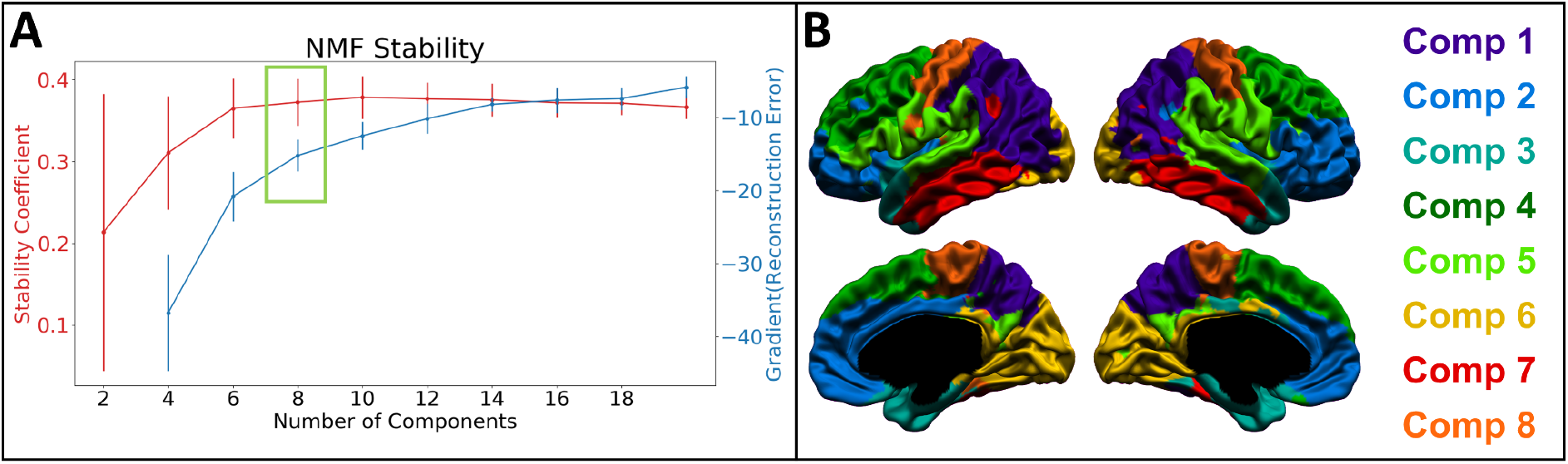
Data-driven parcellation of cortical thickness with NMF. (**A**) For each number of components of the NMF reconstruction, the stability (red) and the gradient of the reconstruction error (blue) are shown. The selected number of components is indicated by the green box. (**B**) Result of the winner-take-all NMF cortical thickness parcellation. Component 1 (purple) represents posterior temporo-parietal regions. Component 2 (blue) represents orbitofrontal, medial-frontal, and anterior cingulate regions. Component 3 (turquoise) includes the medial temporal lobe and part of the temporal pole. Component 4 (dark green) represents posterior frontal regions. Component 5 (light green) represents the superior temporal gyrus and inferior parieto-frontal regions. Component 6 (yellow) represents the occipital lobes. Component 7 (red) represents the inferior and middle temporal gyri. Component 8 (orange) represents the sensorimotor cortex.

### 3.5 Consistent univariate relationships between WMH qT2* signal and clinically-relevant variables

Univariate relationships between WMH measures and different types of clinically-relevant variables were assessed, statistically controlling for age, sex, and years of education. In the PV/deep/SWM parcellation (Fig. 5), we observed significant relationships for WMH volume in global and PV regions with atrophy (medial temporal lobe), cognition (MoCA), clinical group, and cardiovascular risk factors (high BP and cholesterol). Similar relationships were observed for PV and global WMH qT2*, more specifically with atrophy (medial temporal lobe), cognition (MoCA, did not survive FDR correction), clinical group, and cardiovascular risk factors (high cholesterol). While significant group effects were observed, there were no significant pairwise group differences (Supplementary Fig. 2). Other WMH measures did not show robust relationships across all types of clinical variables. When assessing the added value of WMH signal measures above the WMH volume (Supplementary Fig. 3), qT2* remained significantly associated with atrophy (TBV/ICV ratio), clinical group, and cardiovascular risk factors (high BP and high cholesterol), but not cognition.

**Figure 5.**
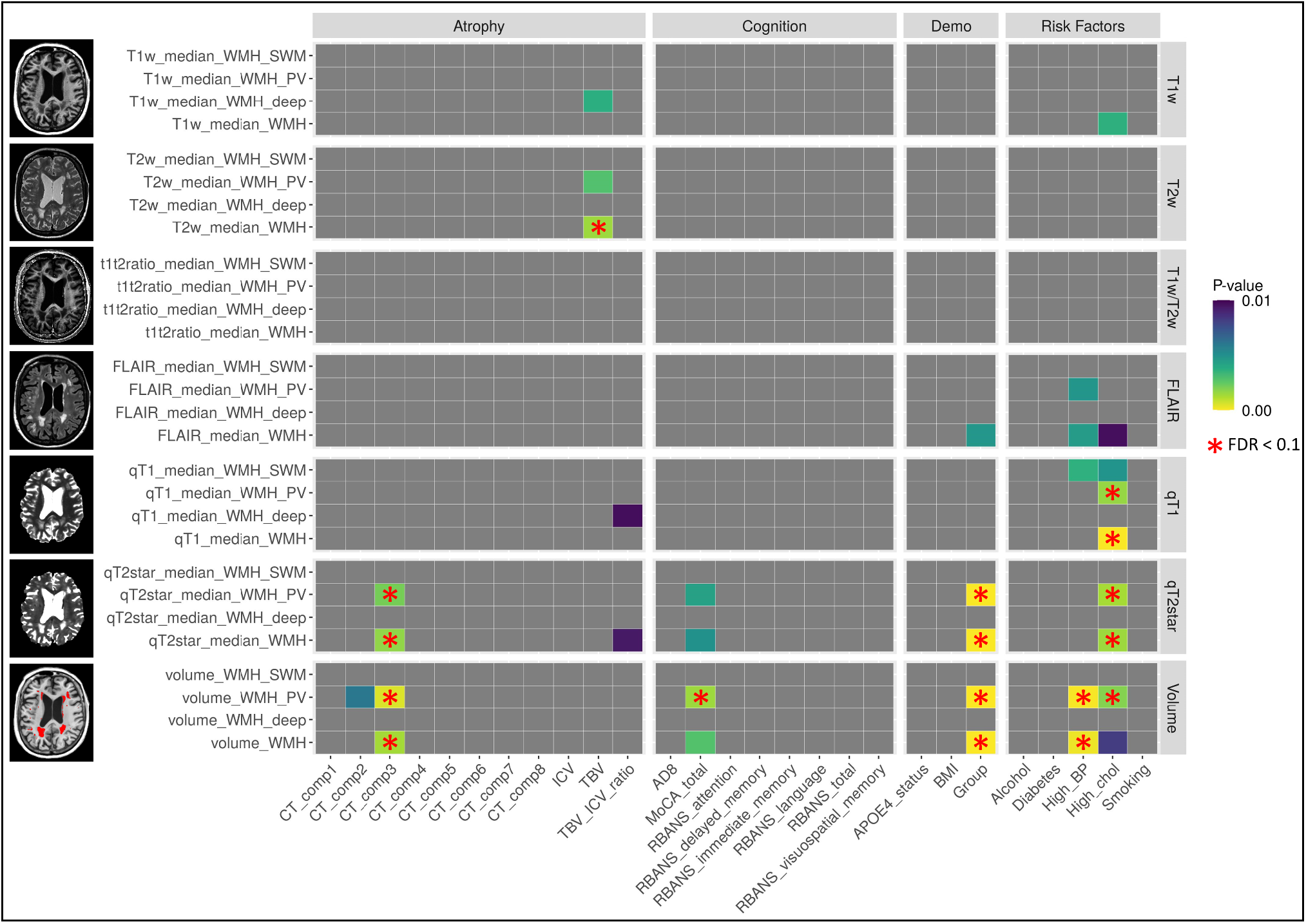
Univariate analyses relating WMH characteristics to clinical variables (PV/deep/SWM parcellation). On the y-axis, all WMH measures (global and parcellated) are grouped by type of image. On the x-axis, all clinical variables are grouped by category. P-values of relationships between each WMH measure and each clinical variable (correcting for age, sex, and education) are shown thresholded at *p*<0.01., with non-significant associations in gray. Yellow colors indicate lower p-values, and purple colors indicate higher p-values. Relationships that survived FDR correction at the 0.1 level are indicated with a red star.

In the lobar parcellation (Supplementary Fig. 4), occipital WMH volume and qT2* were associated with a high number of atrophy, cognitive, and risk factor variables, as well as group effects. WMH volume and qT2* in other lobar regions were also associated, although less extensively, with atrophy (medial temporal lobe), cognition (MoCA), clinical group, and cardiovascular risk factors. Other WMH measures did not show robust relationships across all types of clinical variables.

### 3.6 Multivariate relationship of WMH qT2*, qT1, and FLAIR to age, cognition, and high blood pressure

We further investigated multivariate relationships relating patterns of brain variables (WMH measures and cortical thickness) to patterns of cognitive and demographic variables with PLSC. In the PV/deep/SWM parcellation, LV1 explained the vast majority of the covariance between variables (83%), was significant (*p*=0.0002), and survived split half resampling (Supplementary Fig. 5A). LV3 was also significant (*p*=0.007) and survived split-half resampling but only explained a very small proportion of the covariance (6%), thus is not further analyzed but is available in Supplementary Fig. 5B. In LV1, a pattern of older age, higher AD8, lower immediate memory, attention, global cognitive functioning, and high blood pressure was related to a global pattern of lower cortical thickness and higher WMH burden (Fig. 6). More specifically, the highest contributors to that pattern of brain variables according to the variable-specific bootstrap ratios were decreased cortical thickness in components 5, 3, and 6 (respectively representing superior temporal regions, medial temporal lobe, and occipital regions) and increased WMH volume and qT2* particularly in PV regions. Other WMH signal measures that were significant contributors at the BSR>3.29 level (equivalent to *p*<0.001) were WMH qT2* signal in other white matter regions, qT1 signal in global and SWM WMHs, and FLAIR signal in global and PV WMHs.

**Figure 6.**
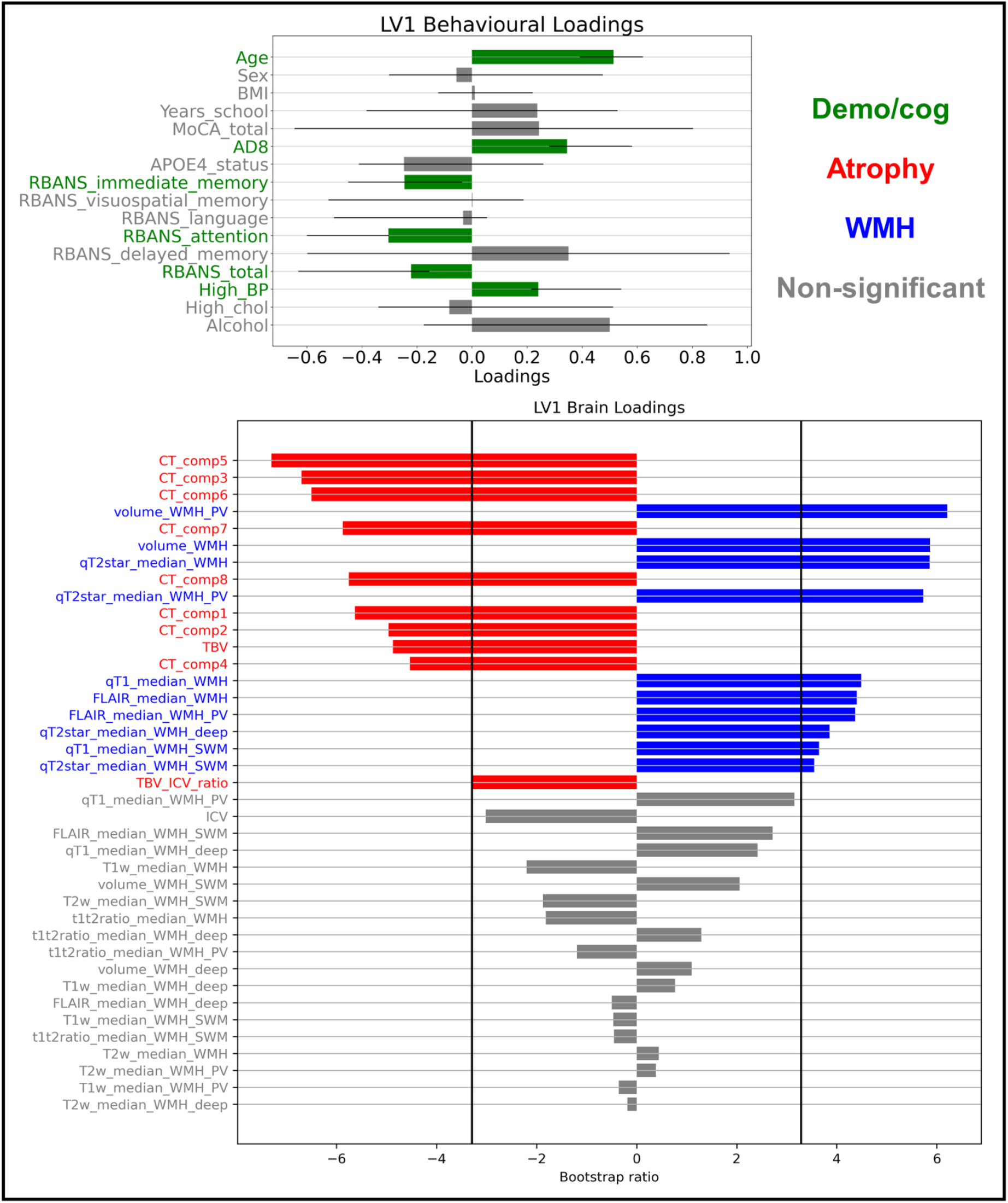
Partial-Least Squares Correlation analysis relating brain variables to cognition and demographics (PV/deep/SWM parcellation). Demographic and cognitive variables are shown in green, atrophy variables in red, WMH variables in blue, and non-significant variables in gray. Top: For each demographic and cognitive variable, the loading on LV1 is proportional to the correlation coefficient on the x-axis. 95% confidence intervals are shown, and variables contribute significantly to the LV (green) if the confidence interval does not cross 0. Bottom: For each brain variable, the bootstrap ratio (BSR) is proportional to the width of the bar on the x-axis. The variables are ordered from top to bottom by BSR value magnitude. Vertical lines at BSR +/-3.29 (equivalent to p<0.001) indicate the significance thresholds.

Similar results were obtained using the lobar parcellation (Supplementary Fig. 6), with the first LV explaining the majority of the variance and showing a multivariate pattern of lower atrophy and higher WMH volume, qT2*, qT1 and FLAIR signal (especially in frontal regions) which is related to a pattern of older age, worse global cognition, high blood-pressure, and lower education level.

## 4. Discussion

### 4.1 qT2* as a potential indicator of tissue damage in WMHs

Our primary finding revealed the qT2* relaxation time of WMHs as a prime candidate for assessing WMH microstructural damage given the differential signal trends it demonstrated relative to NAWM, highlighting a sensitivity to tissue deterioration with age and vascular disease above what would be expected in the global white matter. We further observed consistent associations of WMH qT2* with cortical atrophy, clinical group differences, cardiovascular risk factors, and to a lesser extent cognitive performance. Importantly, qT2* demonstrated predictive clinical value beyond WMH volume, highlighting the potentially complementary nature of these two measures to better characterize WMH severity. Given the known heterogeneity in the microstructural underpinnings of WMHs, qT2* could add crucial clinically-relevant and prognostic information when assessing the overall WMH burden.

Early in the process of WMH genesis, tissue alterations mostly represent accumulations of interstitial water content (i.e., edema) due to disruptions in the blood-brain barrier.^62^ Following tissue damage includes inflammation,^63,64^ demyelination, axonal loss,^15,19,65^ and death of oligodendrocytes.^66^ While a large portion of the qT2* signal can be attributed to variations in interstitial water content (positive association),^67^ studies demonstrated roughly equal contributions of iron (negative association) and myelin (negative association) to the qT2* signal in healthy white matter.^22^ Since most of the magnetic susceptibilities in white matter stems from paramagnetic iron, which is mostly localized in oligodendrocytes,^68^ a reduction in iron and magnetic susceptibilities likely indicates the death of oligodendrocytes. Integrating these known WMH microstructural alterations and sources of qT2* signal, we hypothesize that the increase of qT2* in WMHs could potentially be caused by a combination of increasing interstitial water content and decreasing myelin sheath and oligodendrocyte density. While qT2* might be less specific to any one of these microstructural properties compared to other quantitative MRI measures, it could be sensitive to a unique combination of these properties that best captures the overall severity of WMH tissue damage, thus resulting in the higher clinical associations we observed.

To our knowledge, this is the first study that linked the WMH qT2* signal to clinical variables, although some studies have investigated this measure in a more descriptive manner. Iordanishvili *et al*.^*69*^ observed a significant increase in PV qT2* with WMH volume, an effect that was not observed for NAWM, which supports our observations. One study used myelin water imaging, which separates the quantitative T2 signal into a short (myelin water sensitive) and long component (interstitial water sensitive), to investigate tissue alterations within WMHs.^70^ They observed that the myelin water fraction decreased with WMH volume and was lower in stroke populations, while the intra- and extra-cellular water fraction increased with WMH volume and was not different between stroke and normally-aging populations. Crucially, we extended this body of work by empirically demonstrating associations of the WMH qT2* signal with adverse neurobiological and clinical outcomes.

Interestingly, the clinical associations we observed were mostly for the qT2* signal in PV and occipital WMHs, suggesting that the location of damage is critical. PV WMHs have been more extensively associated with clinical outcomes but tend to represent a lower degree of demyelination compared to other WMHs.^6,16,17^ Similarly, studies show that the clinical impact of WMHs located in posterior regions (i.e., occipital and parietal lobes) is particularly significant in AD.^4,53,71^ As such, microstructural tissue damage of WMHs located in PV and occipital white matter regions, measured by the qT2* signal, could be particularly disruptive for cognitive processes and have a higher clinical impact. Alternatively, qT2* may be particularly sensitive to the tissue damage dynamics of WMHs in those regions.

### 4.2 Limited clinical relevance of other signal measures of WMHs

Other signal measures of WMHs did not meet our criteria for clinical relevance since they did not show consistent associations with all categories of clinical variables and/or did not show divergent signal trends compared to NAWM.

We did not observe significant univariate relationships of WMH qT1 with cognition, atrophy, and demographic variables, but we did observe significant univariate relationships with high cholesterol, as well as significant contributions of WMH qT1 to a multivariate pattern of age and cognition. qT1 in NAWM has been associated mostly with myelin (negative association) to a much larger extent than iron,^22^ and is highly influenced by interstitial water density (positive association).^72^ Interestingly, while the overall qT1 signal was different between WMH and NAWM, we observed that signal trends in aging and disease were highly similar, with signal in both regions slightly increasing with age and WMH volume. This can be explained by one of two possible mechanisms. First, although it is possible that the relationship between qT1 relaxation time and myelin does not hold in WMH tissue, Gouw *et al*.^*18*^ observed that the WMH qT1 signal was independently related to histologically-determined markers of myelin loss, but also axonal loss and microglial activation. The second possible explanation could imply the same rate of demyelination in WMHs and NAWM. Numerous studies have shown that myelin and qT1 alterations are widespread in the NAWM of patients with small vessel disease.^18,62,73,74^ We hypothesize that the initial increase in qT1 is due to edema and that further qT1 increases might indicate a progressive loss of myelin sheaths that occurs at similar rates in WMH and NAWM. Nonetheless, we argue that, based on our results, qT1 is not a sensitive measure of microstructural tissue alterations specific to WMH.

While the WMH FLAIR signal showed highly divergent trends compared to NAWM, we observed few univariate relationships with clinical variables and some contributions to a multivariate pattern of aging and cognition. One study investigated microstructural substrates of FLAIR and T2w signal in WMHs and did not find associations between the intensity (i.e., brightness) of WMHs on these two images and the degree of axonal and myelin degradation.^65^ Since FLAIR, T1w, T2w, and T1w/T2w ratio are qualitative MRI images and are thus more impacted by non-biological sources of variation, it is possible that this added noise compared to quantitative MRI limits the detectable associations with clinical variables and histopathological markers. Nevertheless, qT1 and FLAIR signals could potentially be of value in assessing the WMH microstructural damage given their multivariate relationships with clinical variables. Since FLAIR is not too highly correlated with qT2* signal (especially in PV regions), it could explain different sources of microstructural variations.

The clinical relevance of T1w, T2w, and T1w/T2w ratio measures of WMH severity appears to be limited. Signal trends from all these measures were mostly not significantly different between WMH and NAWM and we observed very few univariate and multivariate relationships with clinical variables. In contrast, some studies have used WMH T1w signal as a measure of WMH severity with the rationale that, since WMHs appear consistently smaller on this type of image, it could potentially be sensitive to more advanced tissue damage and demyelination.^75^ One study reported accelerated WMH T1w signal change after conversion from MCI to dementia, an effect that was not observed for WMH volume.^49^ This discrepancy with our findings could be due to differences in the samples: Dadar *et al*.^*49*^ used a longitudinal sample of 178 MCI subjects who converted to dementia, while our study was cross-sectional and only included 9 participants with dementia. As such, the T1w intensity could be especially sensitive to WMH severity in participants with more advanced neurodegeneration and possibly more advanced small vessel disease. Still, our observations clearly show that other signal measures, such as qT2*, are more sensitive to WMH microstructural tissue damage.

### 4.3 Clinical associations of increased WMH volumes

Our results largely recapitulate previously reported associations of WMH volume with clinical variables. Our observed associations between WMH volume and medial temporal lobe cortical thickness add to the growing body of literature suggesting that WMHs are associated with the stereotypical pattern of neurodegeneration in AD.^9,10,76^ Univariate relationships between WMH volume and cognition were less widespread than previously reported,^6,7^ as associations were restricted to MoCA scores and PV WMH volume, although additional relationships between occipital WMH volume and RBANS subscales (immediate memory, language, and global cognition) were uncovered using a lobar parcellation. While we observed significant overall clinical group differences with respect to WMH volume, there were no significant pairwise group differences in the PV/deep/SWM parcellation. This is possibly due to a lack of power from the small number of AD participants included in the final sample (N=9), which possibly leads to an underestimation of the pathology level in our targeted population. Lastly, our observed associations between WMH volume and hypertension are consistent with the literature.^12,14^

### 4.4 Separating WMHs located in deep and superficial white matter regions

While the classical WMH parcellation only segregates periventricular and deep white matter regions, we differentiated WMHs located in the superficial white matter, which we defined as 1 mm from the cortical gray matter. SWM is mostly composed of association U-fibers connecting cortical gyri and sulci and is thought to be relatively protected from vascular dysfunction since it is doubly vascularized,^77,78^ although WMHs located in SWM regions have been sparsely studied.^49^ However, we observed that there were more WMHs in SWM than in the deep white matter. This is possibly explained by the fact that the total SWM region was the largest on average, representing around 47% of the total white matter in our parcellation scheme (Supplementary Fig. 1). Still, our results do not support reports of vascular protection of SWM since the majority of WMH voxels not in periventricular regions were within 1 mm of cortical gray matter.

### 4.5 Strengths and limitations

The greatest strength of our study is our sample of participants across the AD spectrum, resulting in a higher degree of variability in WMH severity, neurodegeneration, and cognitive functioning, thus increasing statistical power. Our sample is also well-suited for comparing signal measures of WMH since it included five different types of structural MRI acquisitions.

However, our study’s limitations include a somewhat limited sample size to detect associations between brain and clinical variables which are generally of small effect sizes, partly caused by a lack of phenotypical reliability of diagnostic and cognitive measures.^79,80^ This is why we used a stringent criterion to determine clinical relevance (i.e., consistent relationships across types of clinical variables, white matter parcellations, univariate, and multivariate analyses). Despite the moderate sample size, qT2* clearly showed robust associations with neurobiological and clinical variables, and the fact that we observed highly similar clinical effects of WMH volumes with previous findings in the literature brings confidence to the generalizability of our results. Still, smaller associations of other WMH signal measures might not have been detected due to a lack of power.

One technical limitation is that no inhomogeneity corrections for quantitative images were used in this paper. While qT1 maps are not influenced by B1-, proton density (PD), and T2* effects,^81^ several approaches to correct B1+ field inhomogeneities have been proposed^82,83^ but these require additional acquisitions, which we did not have access to. Additionally, while qT2* has been shown to be dependent on the orientation of white matter fibers with respect to the main magnetic field B0,^84,85^ no correction was used in the present paper. Future research investigating the impact of those biases would be necessary to confirm our results and interpretation.

We did not include signal measures from diffusion tensor imaging (DTI), which are traditionally used to assess white matter integrity. Studies show that the DTI-derived microstructure within WMHs scales with WMH volume,^62,70^ and is associated with gait disturbances^86^ and cardiovascular risk factors.^87^ Reports have demonstrated that DTI alterations are detectable before the area appears hyperintense on FLAIR images, thus possibly being sensitive to early tissue alterations of WMHs, and show divergent longitudinal changes compared to NAWM.^88,89^ Hence, DTI metrics also show potential as signal measures of WMH tissue damage severity.

Lastly, it is not clear if our findings are generalizable outside of the AD spectrum. While some studies show generally similar microstructural substrates of WMHs between patients with and without AD,^15^ others report that some WMHs in AD are caused by cortical AD pathology through Wallerian degeneration and not small vessel disease, especially for WMHs located in parietal regions.^19^ However, our sample only contained a small number of AD subjects (*n*=9).

### 4.6 Future directions

Our findings not only highlight that it is possible to characterize the extent of WMH microstructural alterations in vivo but also identify qT2* as the most clinically-relevant MRI marker of tissue damage within WMHs. It will be important in future studies to assess the biological sources of the pathological increase of qT2* in WMHs, for example by relating the WMH qT2* signal to post-mortem histological markers of myelin and iron, as well as immunohistochemistry markers of ischemia, blood-brain barrier dysfunction, and inflammation. Future studies should also assess if the WMH qT2* signal is also clinically-relevant in different populations outside of the AD spectrum with high WMH prevalence, such as pure vascular dementia, psychiatric disorders, and normal pressure hydrocephalus.^16^

An important potential use for assessing the degree of microstructural damage inside WMHs is to determine which WMHs represent irreversible tissue damage (i.e., myelin and axonal loss), or potentially reversible tissue damage (i.e., edema). Indeed, a growing body of literature reports WMH volume reductions in some participants.^90^ Given that the rate of WMH volume change is associated with modifiable cardiovascular risk factors,^90^ it would be particularly relevant to identify if the WMHs of a patient could be reversible following cardiovascular interventions such as hypertensive medication, change in diet, and increased physical activity. WMH signal measures, particularly qT2* but also possibly qT1 and FLAIR, could potentially add critical information in that regard, and as a result, could play a role in personalizing intervention strategies.

## 4.7 Conclusion

In summary, we assessed for the first time the clinical significance of qualitative and quantitative MRI WMH signal measures by relating them to cortical and global atrophy, cognition, clinical group along the AD spectrum, and cardiovascular risk factors. We discovered that qT2* is a relevant marker of WMH microstructural damage, as it showed sensitivity to WMH-specific tissue degradation and was consistently associated with all types of clinical variables across different analysis schemes. qT1 and FLAIR could also be of interest, although to a lesser extent according to our results. We conclude that combining volumetric and signal measures of WMH should be used to improve the characterization of WMH severity in vivo.

