## Supplementary for "Assessment of white matter hyperintensity severity using multimodal MRI in Alzheimer’s Disease"

### Supplementary Methods

#### 1. Quality control

Rigorous manual quality control (QC) was performed for all raw MR images and segmentations. For raw image QC, we excluded participants based on motion artifacts following guidelines established by our group (https://github.com/CoBrALab/documentation/wiki/Motion-Quality-Control-(QC)-Manual).1 We further excluded participants based on inaccuracies in the cortical surface reconstruction (https://github.com/CoBrALab/documentation/wiki/CIVET-Quality-Control-Guidelines) and WMH segmentation errors (i.e., important over- or under-segmentations).

#### 2. MRI sequence parameters

- **T1-weighted**: T1w images were extracted from a magnetization-prepared rapid acquisition gradient echo (MPRAGE) sequence with parameters established by the Alzheimer's Disease Neuroimaging Initiative[2]: repetition time (TR) = 2300 ms; echo time (TE) = 2.98 ms; inversion time (TI) = 900 ms; flip angle (α) = 9º; GRAPPA = 2; slice thickness = 1 mm; 1 mm isotropic in resolution.

- **T2-weighted**: T2w images were extracted from the SPACE sequence with the following parameters: TR = 2500 ms; TE = 198 ms; GRAPPA = 2; slice thickness = 0.64 mm; slice partial Fourier = 6/8; 0.6 mm isotropic in resolution.

- **Fluid Attenuated Inversion Recovery**: The 3D sagittal FLAIR sequence was acquired with the following parameters: TR = 5000 ms; TE = 388 ms; TI = 1800 ms; GRAPPA = 2; slice thickness = 1 mm; 1 mm isotropic in resolution.

- **Quantitative T1**: Quantitative T1 images were extracted from the magnetization-prepared two rapid acquisition gradient echo (MP2RAGE) sequence[3] with the direct output from the Siemens scanner, with the following parameters: TR = 5000 ms; TE = 2.91 ms; TI1 = 700 ms; TI2 = 2500 ms; α1 = 4º; α2 = 5º; GRAPPA = 3; slice thickness = 1 mm; slice partial Fourier = 7/8; 1 mm isotropic in resolution.

- **Quantitative T2***: Quantitative T2* images were extracted from the multi-echo Gradient Recalled Echo (GRE) sequence with 12 echoes with the following parameters: TR = 44 ms; TEs = [2.84, 6.2, 9.56, 12.92, 16.28, 19.64, 23, 26.36, 29.72, 33.08, 36.44, 39.80]; α = 15º; slice thickness = 1 mm; phase partial Fourier = 6/8; 1 mm isotropic in resolution. Specifically, an exponential curve was fit at each voxel using a Python script with the Levenberg-Marquardt curve_fit function[4], and the resulting T2* parameter was extracted with the following expression

$$M = S0 * e(\frac{-t}{T2*})$$

Where M represents the magnetization, S0 represents the maximum signal amplitude which depends on proton density and acquisition parameters, such as the flip angle, t represents the echo time, and T2* represents the time constant.

### 3. Non-negative matrix factorization

To reduce the dimensionality of the cortical thickness data, we used orthonormal-projective non-negative matrix factorization (NMF) to uncover non-overlapping patterns of covariance within the spatially-embedded data, resulting in a data-driven parcellation of the cortical data.[5–7] The code used is publicly available (https://github.com/CoBrALab/cobra-nmf). An input matrix *X* of vertices by subject (*m* x *n*) is decomposed into two matrices: 1) vertices by components representing the spatial parcellation (*m* x *k*; matrix of components *W*) and 2) components by subjects representing the cortical thickness of every subject within each component (*k* x *n*; weight coefficient matrix *H*). The user specifies the number of components k. The decomposition is such that the multiplication of the two output matrices should approximately reconstruct the input matrix. As described by Sotiras *et al.*,[5] the error term the algorithm aims to minimize is shown in Eq. 1.

$$\left|\left|X - WW^tX\right|\right|_F^2 \ where \ WW^t = I \ and \ W \geq 0 \qquad (1)$$

Where || || represents the squared Frobenius norm and *I* represent the identity matrix, which enforces the orthogonality constraint of the columns of $W$. The $W$ matrix is initialized with non-negative double singular value decomposition, which encourages sparsity in the output components and, along with the orthogonality constraint of $W$, allows for the recovery of spatially non-overlapping components through a winner-take-all approach. After the initialization, $W$ is updated using a multiplicative update rule (Eq. 2), where $i$ is the vertex and $j$ is the specific component, and the weight matrix is calculated by projecting the $X$ input matrix onto $W$ (Eq. 3).

$$W'_{ij} = W_{ij} \frac{(XX^tW)_{ij}}{(WW^tXX^tW)_{ij}} \qquad (2)$$

$$H = W^tX \qquad (3)$$

The weight coefficient matrix H is then used as the subject-wise measure of cortical thickness for all analyses.

The input matrix was z-scored across vertices and subjects, and values were shifted by the minimum value to respect the non-negative constraint of NMF. Initialization parameters for the non-negative double singular value decomposition of $W$ were: max iterations = 100,000 and tolerance = 0.00001. The number of components selected was based on the examination of the stability and accuracy of the reconstruction. We first performed a median split of the subjects based on age, then 10 subsets in each age group were selected at random. At each specified number of components (between 2 and 20), and for each of the 10 subsets of participants, we assessed the cosine similarity between the NMF reconstructions of the younger and older group, representing the stability of the spatial components. During this process, we also calculated the gradient of the reconstruction error at each granularity to quantify the accuracy of the reconstructions. Finally, we chose the final number of components that we subsequently used for all analyses of CT to respect the principle of parsimony (i.e., the smallest number of components with appropriate accuracy and stability).

### 4. Partial-Least Squares Correlation

In order to investigate how signal measures of WMH help predict demographics and behaviors in conjunction with traditional brain markers (i.e., cortical thickness and WMH volume), we used the multivariate technique Partial Least Squares Correlation (PLSC). To proceed, we used behavioral PLSC correlation with the Python package pyls (https://github.com/rmarkello/pyls) with Python version 3.9.7 and scikit-learn version 1.0.2, which performed singular vector decomposition on a correlation matrix of every brain variable to every cognitive and demographic variable.[8–11] This results in uncorrelated latent variables (LV) representing linear combinations of cognitive and demographic variables that maximally covary with linear combinations of brain variables. We assessed the significance of each LV by performing PLSC with 5,000 permutations on the brain data matrix, resulting in a null distribution of singular values from which we derive a non-parametric p-value for each LV. We further assessed the reliability of each LV with split-half analysis.[12] Finally, bootstrap resampling is used to examine the significance of each individual predictor to each LV. We generated 5,000 random samplings (with replacement) of both the brain and cognition/demographics data, from which we extracted a distribution of singular vector weights for each individual predictor. The resulting distribution is used to calculate 95% confidence intervals for cognitive/demographic variables and calculate the bootstrap ratio (BSR; dividing the singular vector weight by the standard error) for brain variables. We used a threshold of BSR=3.29 to determine if predictors significantly contributed to the observed patterns, equivalent to a p-value of 0.001. Of note, a minimum amount of variance for each variable is required in PLSC and most cardiovascular risk factor variables did not have enough variance. Hence, we removed diabetes and smoking history variables from this analysis (only keeping alcohol consumption, high blood pressure, and high cholesterol variables).

### Supplementary Figures

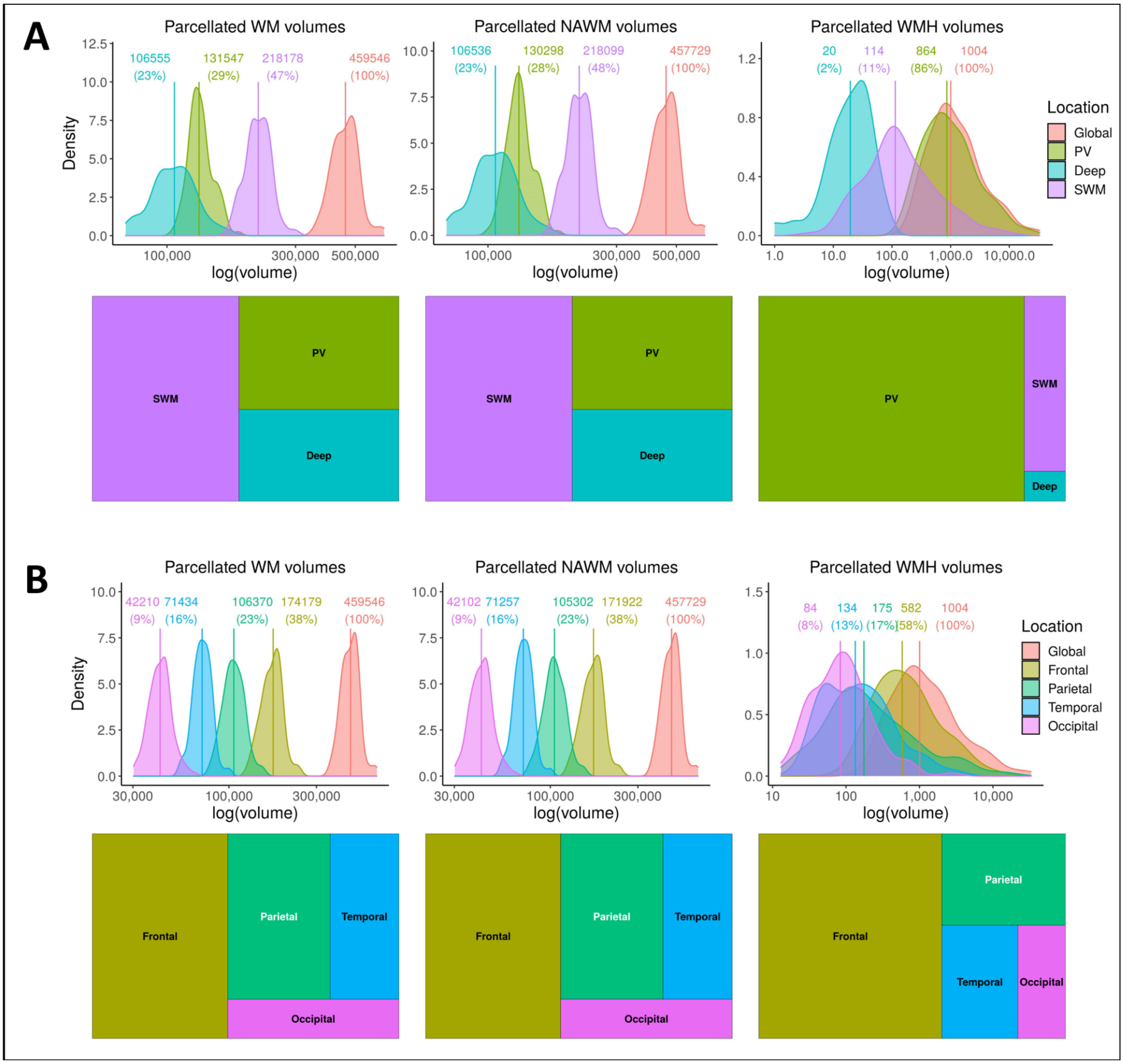


**Supplementary Figure 1 Results of white matter parcellations.** (**A**) Results of non-imputed periventricular/deep/superficial white matter parcellation. (**B**) Results of lobar parcellation. **Top**: Density plots of the global and parcellated volumes of WM, NAWM, and WMH, using a log scale for the volume on the x-axis. Vertical lines represent the median volume. The associated value of the median volume and proportion relative to the global volume are shown. **Bottom**: Proportions of each parcellated volume relative to the global volume are visualized with a tree map.

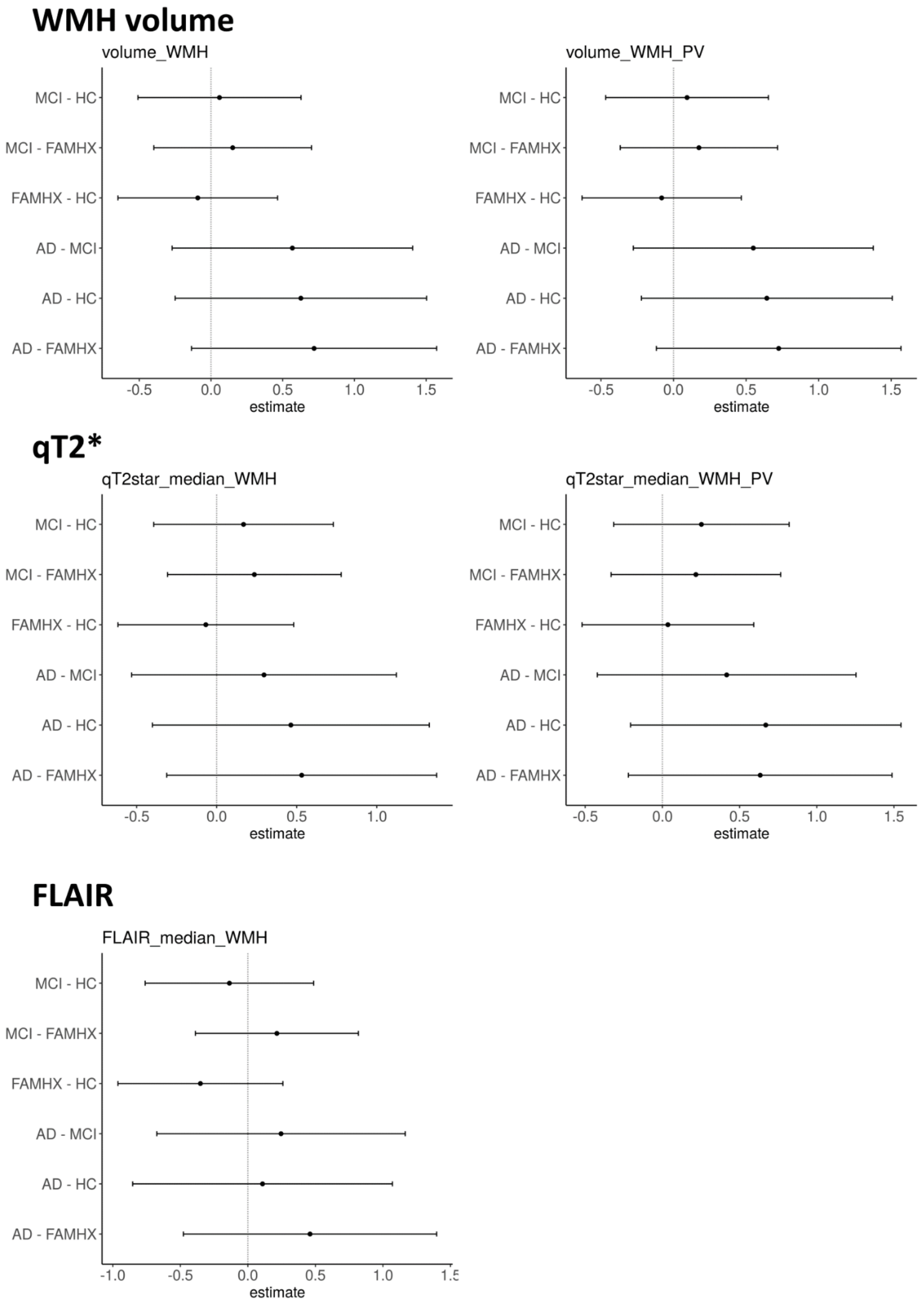


**Supplementary Figure 2 Univariate analyses relating WMH characteristics to clinical variables (PV/deep/SWM parcellation): Pair-wise group differences.** All possible pairwise group differences were assessed with Tukey comparisons for significant overall group effects previously detected. Each group comparison is indicated on the y-axis, the estimate is indicated by the point on the x-axis, and 95% confidence intervals are shown.

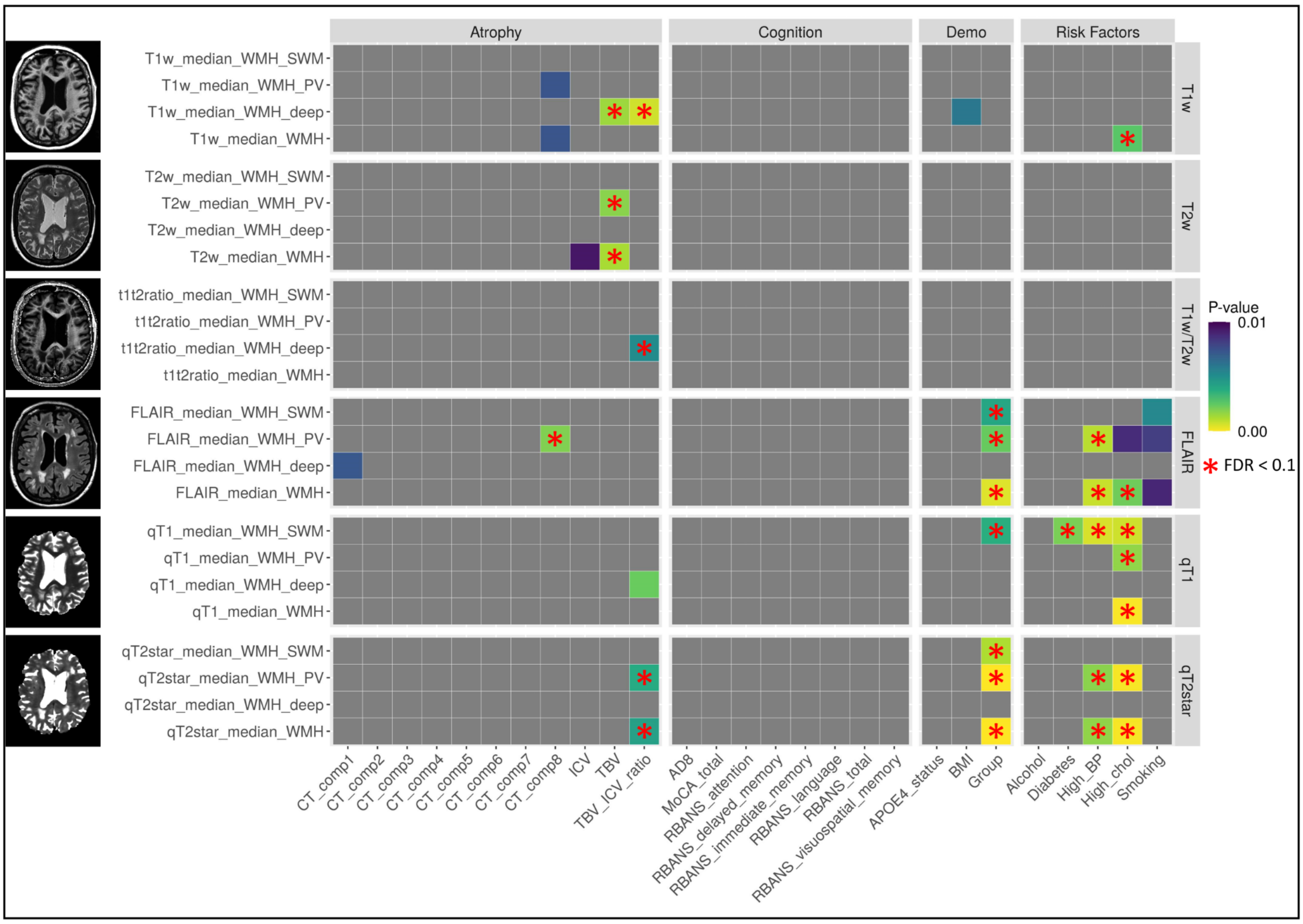


**Supplementary Figure 3 Univariate analyses relating WMH signal measures to clinical variables (PV/deep/SWM parcellation): predictive value above WMH volume.** On the y-axis, all WMH measures (global and parcellated) are grouped by type. On the x-axis, all clinical variables are grouped by type. P-values of relationships between each WMH measure and each clinical variable (correcting for age, sex, education, and region-specific WMH volume) are shown thresholded at $p<0.01$, with non-significant associations in gray. Yellow colors indicate smaller p-values, and purple colors indicate bigger p-values. Relationships that survived FDR correction at the 0.1 level are indicated with a red star.

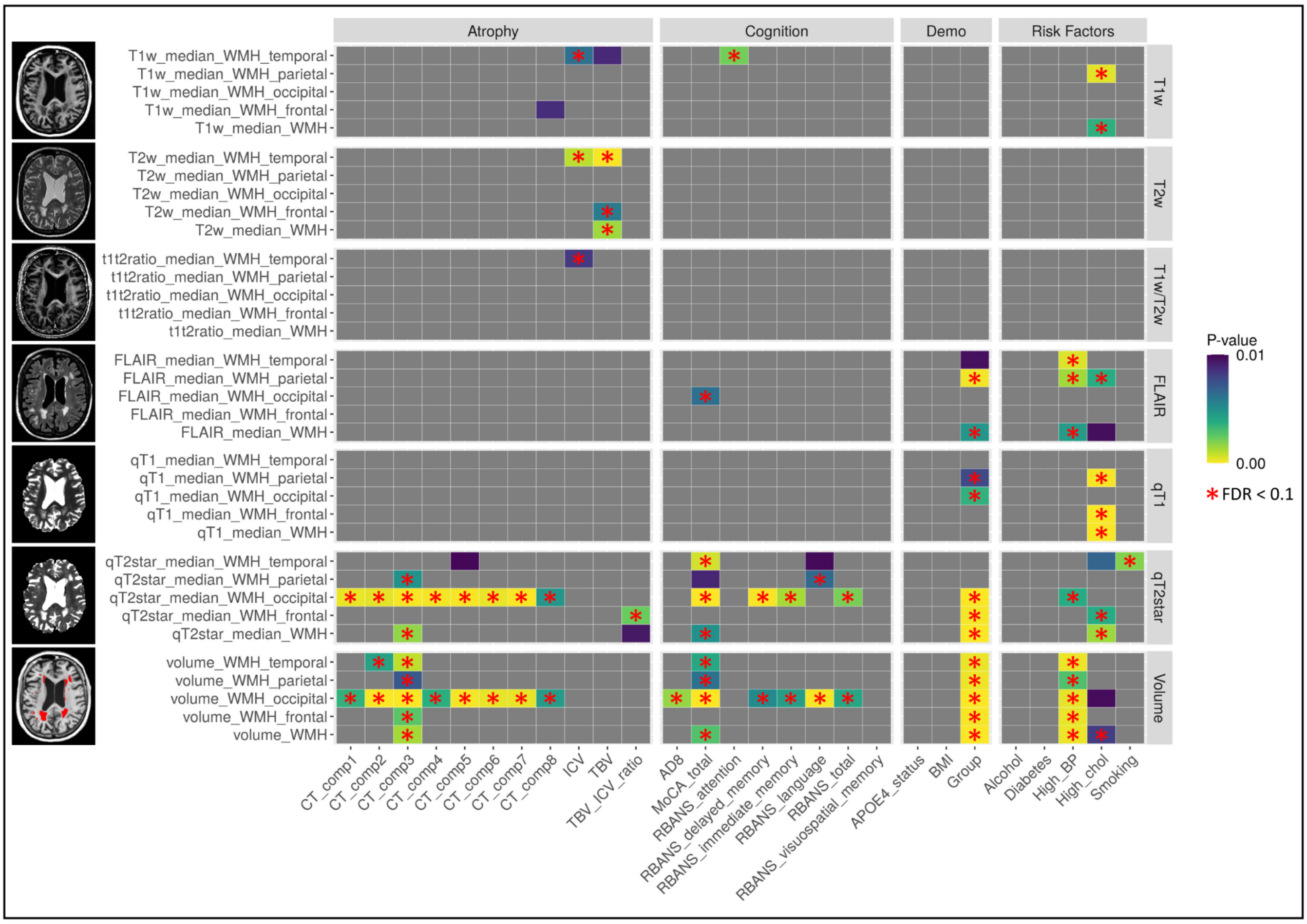


**Supplementary Figure 4 Univariate analyses relating WMH characteristics to clinical variables (lobar parcellation).** On the y-axis, all WMH measures (global and parcellated) are grouped by type. On the x-axis, all clinical variables are grouped by type. P-values of relationships between each WMH measure and each clinical variable (correcting for age, sex, and education) are shown thresholded at $p<0.01$, with non-significant associations in gray. Yellow colors indicate smaller p-values, and purple colors indicate bigger p-values. Relationships that survived FDR correction at the 0.1 level are indicated with a red star.

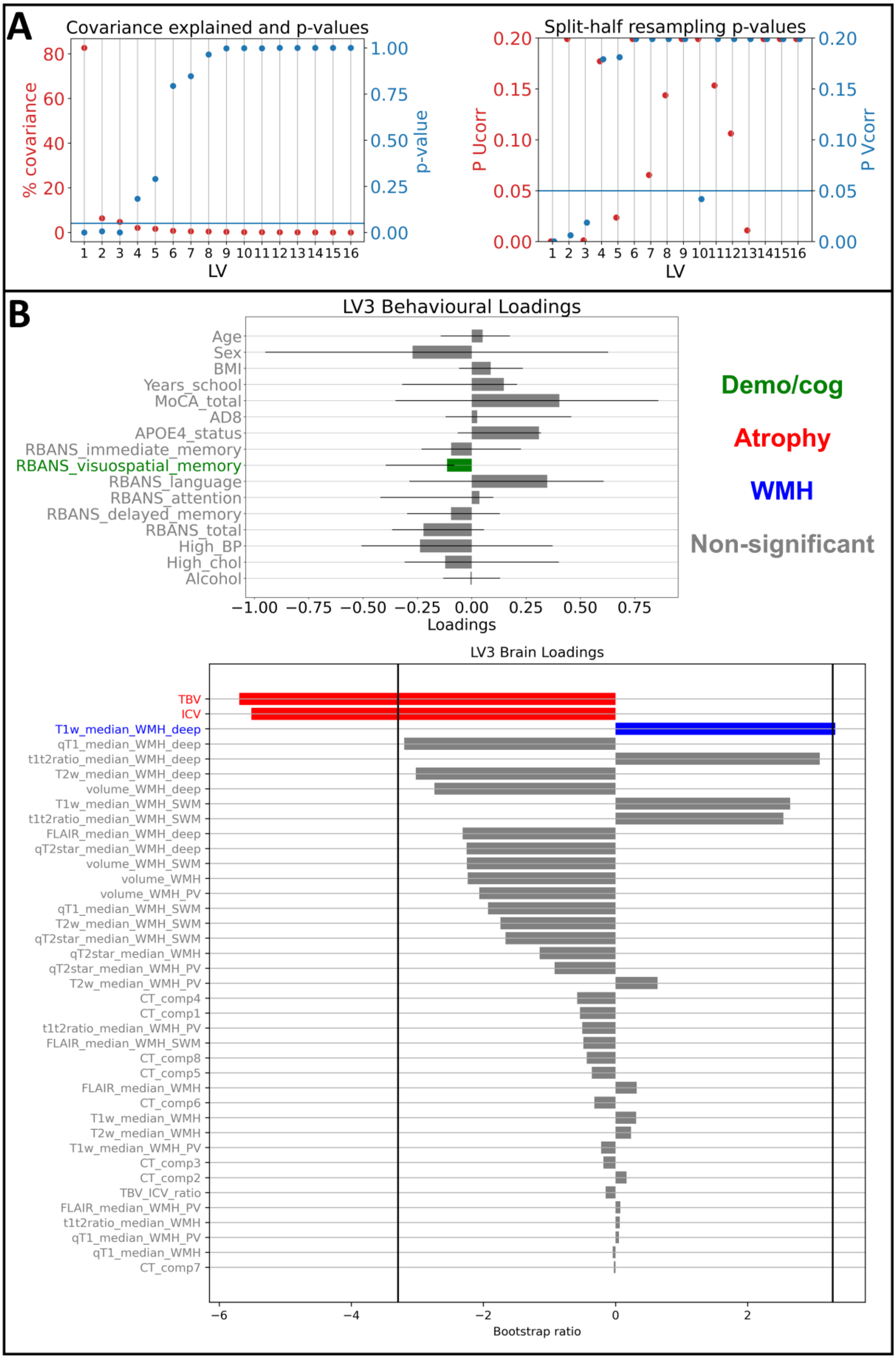

A
Covariance explained and p-values
% covariance
p-value
LV
Split-half resampling p-values
P Ucorr
P Vcorr
B
LV3 Behavioural Loadings
Age
Sex
BMI
Years_school
MoCA_total
AD8
APOE4_status
RBANS_immediate_memory
RBANS_visuospatial_memory
RBANS_language
RBANS_attention
RBANS_delayed_memory
RBANS_total
High_BP
High_chol
Alcohol
Loadings
Demo/cog
Atrophy
WMH
Non-significant
LV3 Brain Loadings
TBV
ICV
T1w_median_WMH_deep
qT1_median_WMH_deep
t1t2ratio_median_WMH_deep
T2w_median_WMH_deep
volume_WMH_deep
T1w_median_WMH_SWM
t1t2ratio_median_WMH_SWM
FLAIR_median_WMH_deep
qT2star_median_WMH_deep
volume_WMH_SWM
volume_WMH
volume_WMH_PV
qT1_median_WMH_SWM
T2w_median_WMH_SWM
qT2star_median_WMH_SWM
qT2star_median_WMH
qT2star_median_WMH_PV
T2w_median_WMH_PV
CT_comp4
CT_comp1
t1t2ratio_median_WMH_PV
FLAIR_median_WMH_SWM
CT_comp8
CT_comp5
FLAIR_median_WMH
CT_comp6
T1w_median_WMH
T2w_median_WMH
T1w_median_WMH_PV
CT_comp3
CT_comp2
TBV_ICV_ratio
FLAIR_median_WMH_PV
t1t2ratio_median_WMH
qT1_median_WMH_PV
qT1_median_WMH
CT_comp7
Bootstrap ratio

**Supplementary Figure 5 Partial-Least Squares Correlation analysis relating brain to cognition and demographics (PV/deep/SWM parcellation): significance, effect size, reliability, and 3rd latent variable.** (**A**) **Left**: Covariance explained (red) and significance (blue) of each latent variable. The horizontal blue line represents the significance threshold of p-values at 0.05. **Right**: Split-half analysis of the reliability of each latent variable. P-values of the correlations of the left (Ucorr; red) and right (Vcorr; blue) singular vectors are shown. The horizontal blue line represents the significance threshold of p-values at 0.05, and p-values from both the left and right singular vectors need to be below 0.05 for the LV to be considered reliable. (**B**) Third latent variable, which is significant and survived split-half analysis. Demographic and cognitive variables are indicated in green, atrophy variables are indicated in red, WMH variables are indicated in blue, and non-significant variables are indicated in gray. **Top**: For each demographic and cognitive variable, the loading on LV3 is proportional to the width of the bar on the x-axis. 95% confidence intervals are shown, and variables have non-significant contributions to the LV if the confidence interval crosses 0. **Bottom**: For each brain variable, the bootstrap ratio (BSR) is proportional to the width of the bar on the x-axis. The variables are ordered from top to bottom with the absolute BSR values. Vertical lines at BSR +/- 3.29 (equivalent to $p<0.001$) indicate the significance thresholds.

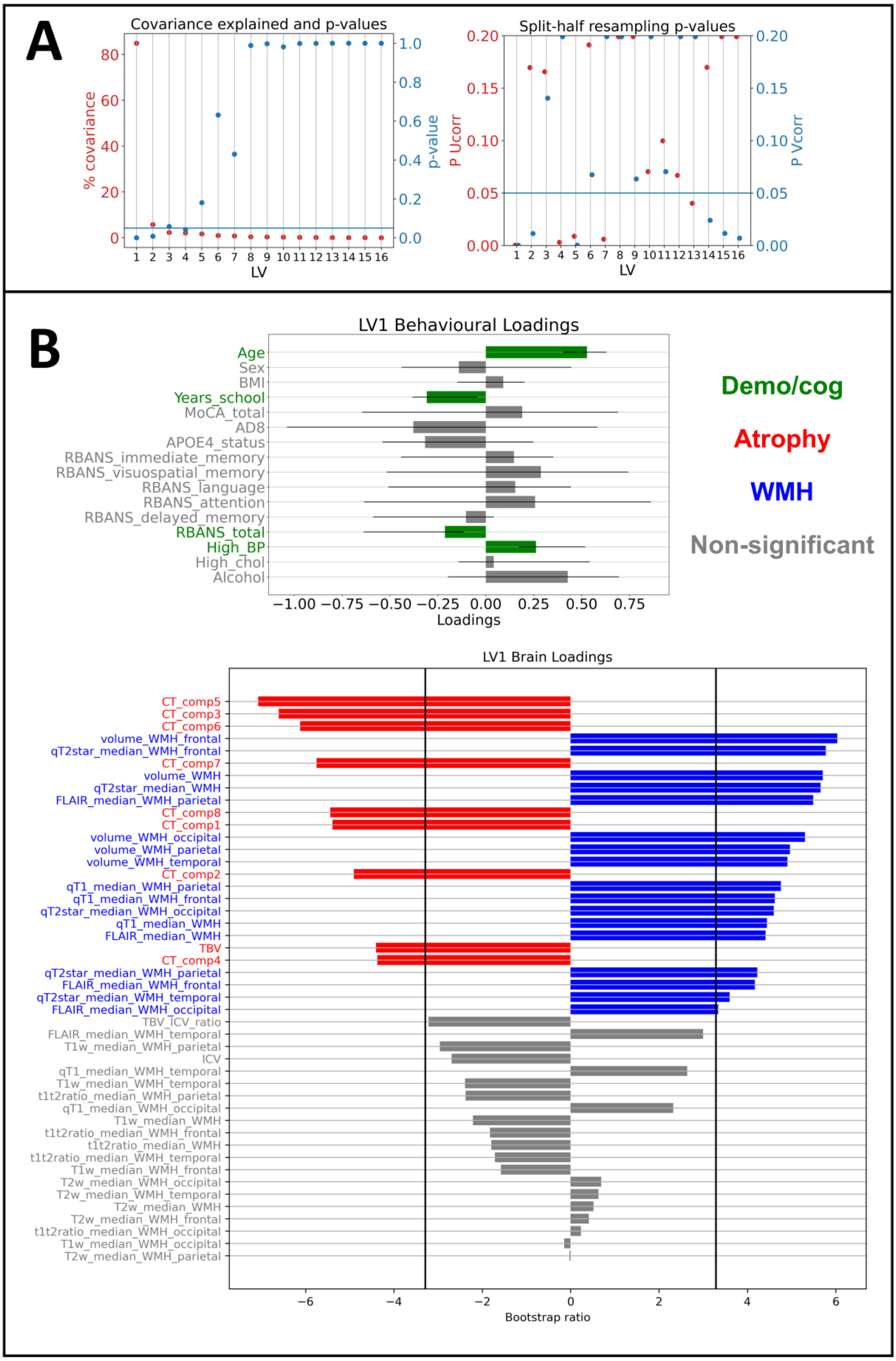
A
Covariance explained and p-values
% covariance
p-value
LV
Split-half resampling p-values
P Ucorr
P Vcorr
B
LV1 Behavioural Loadings
Age
Sex
BMI
Years_school
MoCA_total
AD8
APOE4_status
RBANS_immediate_memory
RBANS_visuospatial_memory
RBANS_language
RBANS_attention
RBANS_delayed_memory
RBANS_total
High_BP
High_chol
Alcohol
Loadings
Demo/cog
Atrophy
WMH
Non-significant
LV1 Brain Loadings
CT_comp5
CT_comp3
CT_comp6
volume_WMH_frontal
qT2star_median_WMH_frontal
CT_comp7
volume_WMH
qT2star_median_WMH
FLAIR_median_WMH_parietal
CT_comp8
CT_comp1
volume_WMH_occipital
volume_WMH_parietal
volume_WMH_temporal
CT_comp2
qT1_median_WMH_parietal
qT1_median_WMH_frontal
qT2star_median_WMH_occipital
qT1_median_WMH
FLAIR_median_WMH
TBV
CT_comp4
qT2star_median_WMH_parietal
FLAIR_median_WMH_frontal
qT2star_median_WMH_temporal
FLAIR_median_WMH_occipital
TBV_ICV_ratio
FLAIR_median_WMH_temporal
T1w_median_WMH_parietal
ICV
qT1_median_WMH_temporal
T1w_median_WMH_temporal
t1t2ratio_median_WMH_parietal
qT1_median_WMH_occipital
T1w_median_WMH
t1t2ratio_median_WMH_frontal
t1t2ratio_median_WMH
t1t2ratio_median_WMH_temporal
T1w_median_WMH_frontal
T2w_median_WMH_occipital
T2w_median_WMH_temporal
T2w_median_WMH
T2w_median_WMH_frontal
t1t2ratio_median_WMH_occipital
T1w_median_WMH_occipital
T2w_median_WMH_parietal
Bootstrap ratio

**Supplementary Figure 6 Partial-Least Squares Correlation analysis relating brain to cognition and demographics (lobar parcellation): significance, effect size, reliability, and 1st latent variable.** (**A**) **Left**: Covariance explained (red) and significance (blue) of each latent variable. The horizontal blue line represents the significance threshold of p-values at 0.05. **Right**: Split-half analysis of the reliability of each latent variable. P-values of the correlations of the left (Ucorr; red) and right (Vcorr; blue) singular vectors are shown. The horizontal blue line represents the significance threshold of p-values at 0.05, and p-values from both the left and right singular vectors need to be below 0.05 for the LV to be considered reliable. (**B**) First latent variable, which is significant and survived split-half analysis. Demographic and cognitive variables are indicated in green, atrophy variables are indicated in red, WMH variables are indicated in blue, and non-significant variables are indicated in gray. **Top**: For each demographic and cognitive variable, the loading on LV1 is proportional to the width of the bar on the x-axis. 95% confidence intervals are shown, and variables have non-significant contributions to the LV if the confidence interval crosses 0. **Bottom**: For each brain variable, the bootstrap ratio (BSR)is proportional to the width of the bar on the x-axis. The variables are ordered from top to bottom with the absolute BSR values. Vertical lines at BSR +/- 3.29 (equivalent to $p<0.001$) indicate the significance thresholds.
